# Taxonomy of the apicomplexan symbionts of coral, including Corallicolida ord. nov., reassignment of the genus *Gemmocystis,* and description of new species *Corallicola aquarius* gen. nov. sp. nov. and *Anthozoaphila gnarlus* gen. nov. sp. nov

**DOI:** 10.1101/2020.11.23.395046

**Authors:** Waldan K. Kwong, Nicholas A.T. Irwin, Varsha Mathur, Ina Na, Noriko Okomoto, Mark J.A. Vermeij, Patrick J. Keeling

## Abstract

Corals (Metazoa; Cnidaria; Anthozoa) have recently been shown to play host to a widespread and diverse group of intracellular symbionts of the phylum Apicomplexa. These symbionts, colloquially called ‘corallicolids’, are mostly known through molecular analyses, and no formal taxonomy has been proposed. Another apicomplexan, *Gemmocystis cylindrus* (described from the coral *Dendrogyra cylindrus*), may be related to corallicolids, but lacks molecular data. Here, we isolate and describe motile trophozoite (feeding) corallicolids cells using microscopic (light, SEM, and TEM) and molecular phylogenetic analysis to provide the basis for a formal description. Phylogenetic analyses using nuclear and plastid rRNA operons, and three mitochondrial protein sequences derived from single cell transcriptomes, all confirm that these organisms fall into a discrete deep-branching clade within the Apicomplexa not closely related to any known species or major subgroup. As a result, we assign this clade to a new order, Corallicolida ord. nov., and family, Corallicolidae fam. nov. We describe a type species, *Corallicola aquarius* gen. nov. sp. nov. from its *Rhodactis* sp. host, and also describe a second species, *Anthozoaphila gnarlus* gen. nov. sp. nov., from the coral host *Madracis mirabilis*. Finally, we propose reassigning the incertae sedis taxon *G. cylindrus* from the order Agamococcidiorida to the Corallicolida, based on similarities in morphology and host localization to that of the corallicolids.

---

The phylum Apicomplexa is a diverse lineage of intracellular symbionts of animals, that infect cells of virtually all animal species that have been examined. In addition to well studied vertebrate hosts, such as humans (e.g., *Toxoplasma, Plasmodium*), cattle and chicken (*Eimeria*), and dogs (*Hepatozoon*), a large number of invertebrates are also known to harbour apicomplexans (Rueckert 2019). While commonly classified as obligate parasites, apicomplexans can in reality engage in a range of interactions with their hosts, including mutualism and commensalism (Rueckert 2019; Paight et al. 2019). Apicomplexans evolved from phototrophic ancestors, retain a relict non-photosynthetic plastid, and are closely related to a collection of free-living predators and two species of photosynthetic algae that are associated with coral reef environments (McFadden and Waller 2005; Moore et al. 2008; Janouškovec et al. 2010; Mathur et al. 2018). Recent studies have identified another link with corals, in this case stemming from environmental sequence data that revealed the presence of an apicomplexan lineage that is prevalent in corals worldwide (Janouškovec et al. 2012; Mathur et al. 2018; Kwong et al. 2019; Vohsen et al. 2020). Members of this lineage, informally named ‘corallicolids’, exhibit substantial sequence diversity and variation in host association patterns, and their phylogenetic position, while not completely resolved, indicates that they do not fall within any of the currently recognized subgroups within apicomplexan taxonomy (Janouškovec et al. 2012; Kwong et al. 2019; Vohsen et al. 2020).

Despite a growing understanding of the diversity and distribution of this lineage, there remains no formal classification of it or its members. This is largely due to previous work generating non-comparable types of data from different potential members of the group. A coral-associated apicomplexan was first formally described by Upton and Peters (Upton and Peters 1986), based on morphology from histological sections of coral tissues. This organism, named *Gemmocystis cylindrus*, was proposed to be a member of the Agamococcidia. However, the description was not associated with any molecular data and no DNA sequence data has ever been procured from a verified sample of *G. cylindrus*. Later, an apicomplexan 18S rRNA gene from coral was identified by Toller et al. (Toller et al. 2002), and given the designation “genotype N”. These sequences were found to branch sister to the Eucoccidia clade (containing members such as *Toxoplasma* and *Eimeria*), but these sequences were never linked to a visualized cell or any other data. Subsequently, a lineage of apicomplexan plastid-encoded 16S rRNA gene sequences were also discovered to be associated with corals and designated “ARL-V” (Janouškovec et al. 2012). These were found to branch sister to all other apicomplexan plastids, but again were not linked to a specific cell or other data.

The taxonomic conclusions from these three kinds of data were not only contradictory, but because the data were not directly comparable, it remained uncertain if these represented one or more than one lineage of apicomplexan. This was ultimately reconciled through the combined use of microscopy and metagenomic sequencing from both plastid and nuclear genomes, leading to the identification of a single clade of coral-infecting apicomlexans, informally dubbed “corallicolids” (Kwong et al. 2019). Here, we isolate and document the first living cells from this clade, use single cell transcriptome data to confirm their identity, and propose a formal taxonomic description of this lineage, along with two new species, to serve as points-of-reference for future studies of this group.

## MATERIAL AND METHODS

### Sample collection and light microscopy

Corallicolid samples from the green mushroom coral, *Rhodactis* sp., were taken from an aquarium in Vancouver, BC, Canada, as described in a previous study (Kwong et al. 2019). Based on morphology and the COX1 sequence (MH320096), the host specimen is part of the *Rhodactis inchoate* / *Rhodactis indosinensis* clade, a coralimorph native to the Indo-Pacific (Oh et al. 2019). Two other corallicolids (sample_10 and sample_27) were isolated from anthozoans collected from Curaçao in April 2019, under the permits of the Dutch Antillean Government (Government reference: 2012/48584) provided to the CARMABI Foundation (CITES Institution code AN001). Fragments of the yellow pencil coral *Madracis mirabilis* (sample_10) and the golden zoanthid *Parazoanthus swiftii* (sample_27) were crushed with a mortar and pestle, then viewed under a Leica DM IL LED inverted microscope. Video was recorded using an attached Sony A6000 digital camera. Individual cells with apicomplexan-like sporozoite morphology were isolated using glass micropipettes, washed twice in 0.2 μm filtered seawater, and were transferred into tubes containing 3 μl of RNase inhibitor solution (1 part 20 U μl^−1^ RNase Inhibitor [Thermo Fisher Scientific] in 19 parts 0.2% Triton X-100) and then frozen. Micrographs of the cells used for transcriptome sequencing are shown in Fig. S1.

### Transcriptome sequencing

Generation of cDNA via reverse transcription was carried out using the Smart-Seq2 protocol (Picelli et al. 2014), and cDNA concentration was quantified with a Qubit 2.0 Fluorometer (Thermo Fisher Scientific). Sequencing libraries were then prepared using the Nextera XT protocol (Illumina). Sample_10 was sequenced on the Illumina NextSeq platform using 2 × 150 bp paired-end reads, and sample_27 was sequenced on the Illumina MiSeq v2 platform using 2 × 250 bp paired-end reads. The sequencing reads were trimmed using TrimGalore 0.6.0 (Martin 2011) to remove adapter and primer sequences, and assembled with rnaSPAdes 3.13.2 (Bushmanova et al. 2019).

### Phylogenetic tree building

Nuclear 18S and 28S rRNA genes, mitochondrial genes (*cox1*, *cox3*, and *cob*), and plastid 16S and 23S rRNA genes were retrieved from the assembled transcripts based on BLAST similarity to existing sequences. Previously published sequences were retrieved from NCBI GenBank and by BLAST searches of genomic and transcriptomic datasets (Table S1). To maximize phylogenetic resolution for corallicolids, 18S rRNA gene sequences < 800 bp in length were excluded; for the 16S rRNA gene, sequences < 1000 bp were excluded. Ribosomal RNA genes were predicted using RNAmmer 1.2 (Lagesen et al. 2007) and aligned using MUSCLE (Edgar 2004). Mitochondrial proteins were aligned with MAFFT (Katoh et al. 2002). Sequences were trimmed to remove unaligned 5’ and 3’ ends and then concatenated. Maximum likelihood trees were built in IQ-TREE 1.5.4 (Nguyen et al. 2015) with 1000 bootstrap replicates. Substitution models were selected using ModelFinder (Kalyaanamoorthy et al. 2017). Trees inferred using the Bayesian method were built in MrBayes 3.2.6 (Ronquist et al. 2012), using 10^6^ generations, sampling every 500 generations, and a burn-in of 0.25. A haplotype network of the 16S rRNA gene was built with PopART (Leigh and Bryant 2015), using the median joining algorithm (Bandelt et al. 1999).

### Electron Microscopy

For scanning electron microscopy, crushed coral slurries were fixed by mixing with 2.5% glutaraldehyde in seawater and kept at 4°C. Samples were then rinsed 3 × with seawater and postfixed with 1% osmium tetroxide in 0.1 M sodium cacodylate pH 7.34. Samples were washed 3 × with 0.1 M sodium cacodylate and placed on a coverslip coated with 0.1% poly-L-lysine, then rinsed with distilled water and air dried in a humidified chamber. Dehydration was carried out with successive 30%, 50%, 70%, 80%, 90%, 95%, and 3 × 100% ethanol washes. Samples were dried with a Tousimis Autosamdri 815B critical point dryer, and sputter coated using a Cressington 208HR high resolution sputter coater. Samples were viewed with a Hitachi S-2600 variable pressure SEM. Transmission electron microscopy was conducted as previously described (Kwong et al. 2019).

### Data availability

Mitochondrial and rRNA gene sequences generated in this study are deposited in NCBI GenBank under accession numbers MW192638–MW192642 and MW206707–MW206712. Accession numbers of sequences used to build phylogenies are listed in Table S1.

## RESULTS AND DISCUSSION

### Description

The mesenterial filaments of the green mushroom coral *Rhodactis* sp. has previously been shown by TEM and in-situ hybridization to be infected with a corallicolid (Kwong et al. 2019). We re-examined this tissue from a specimen kept in an aquarium by crushing filaments and examining them under light microscopy. Extracellular, motile cells with morphology reminiscent of apicomplexan motile or trophic stages stages were observed, albeit rarely. These cells were no more than 15 μm in length and 5 μm in width and exhibited adhesion to the substrate (Fig. 1C). Tissue sections were viewed under transmission electron microscopy, which showed the presence of intracellular cells displaying apicomplexan ultrastructural features (Fig. 1A, B). These include a subpellicular microtubule network, mitochondria with tubular cristae, and micronemes located towards the apical end of the cells. Numerous, distinctive dark-staining organelles up to 1.6 μm in length were also observed. Cells were encased within a parasitophorous vacuole, and two or more cells were often found within the same vacuole. This is suggestive of a reproductive process, plausibly endodyogeny since they did not display the rigid outer wall that is characteristic of oocysts and tissue cysts (Ferguson and Dubremetz 2007). However, the instance of two daughter cells developing within a mother cell (a direct feature of endodyogeny) was not clearly observed, and hence this could also represent another process. We propose the name *Corallicola aquarius* gen. nov., sp. nov. for this corallicolid lineage hosted by *Rhodactis* sp.

**Figure 1.**
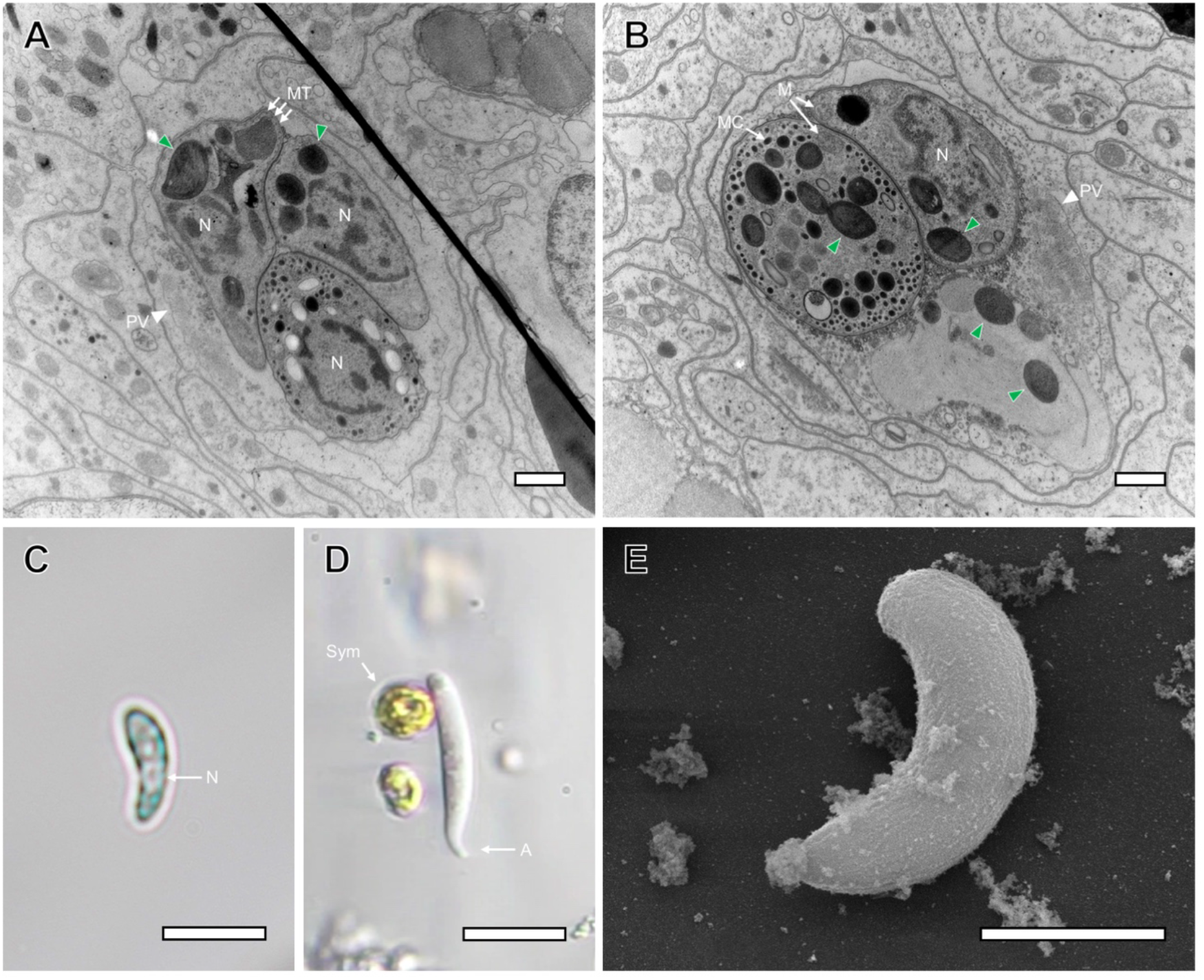
Morphology and structure of corallicolid cells. **A., B.** Transmission electron micrographs of *C. aquarius* from *Rhodactis* sp. These cells may represent endodyogeny or another reproductive life stage. Scale bar = 1 μm. In **A.**, three cells are present, within a putative parasitophorous vacuole (PV). In **B.**, two cells are visible, adjacent to remnant cellular material from a possible third cell. Distinctive, darkly staining organelles (examples indicated with green arrows) are found throughout, including in the remains of the third cell in **B**. Mitochondria, M; micronemes, MC; microtubule network, MT. **C.** Light micrograph of *C. aquarius* from the green mushroom coral (*Rhodactis* sp.). Nucleus, N. Scale bar = 10 μm. **D.** Light micrograph of *A. gnarlus* from the yellow pencil coral (*M. mirabilis*), alongside cells of Symbiodiniaceae (Sym) from the same sample. Anterior rostrum, A. Scale bar = 10 μm. **E.** Scanning electron micrograph of *A. gnarlus* from coral sample_10 (*M. mirabilis*). Scale bar = 5 μm. Cells in **C.–E.** may represent sporozoites or other motile life stage.

Samples of the yellow pencil coral, *M. mirabilis,* from Curaçao (southern Caribbean) were also found to contain cells with apicomplexan-like morphologies, resembling that of sporozoites or a comparable motile life stage (Fig. 1D, E). Cells were modestly elastic in size and shape, but were no more than 20 μm in length and 5 μm in width. The cells displayed gliding motility, with movement directed towards the anterior and generally following a circular, clockwise path of travel (Video S1, Video S2). A flexible rostrum formed the anterior end, while the posterior end was characterized by strong affinity for adhesion to the substrate. These observations are consistent with the typical actin-myosin based gliding locomotion of the Apicomplexa (Frénal et al. 2017). Oocysts or other obvious reproductive forms were not observed in these samples. We propose the name *Anthozoaphila gnarlus* gen. nov., sp. nov. for the cells collected from *M. mirabilis*. A very similar motile cell was found in a distantly-related host, the golden zoanthid, *P. swiftii* (Fig. S1), which was subsequently confirmed by molecular data to be extremely closely related to *Anthozoaphila gnarlus* (see below).

### Taxonomic assignment

We performed RNA sequencing on several single cells of *A. gnarlus*, isolated from *M. mirabilis.* While none of these provided sufficient depth of coverage to be considered complete transcriptomes, we were able to retrieve transcripts corresponding to several nuclear, mitochondrial, and plastid genes. We also identified the nuclear rRNA operon and all three mitochondrial protein-coding genes from the *A. gnarlus*-like symbiont isolated from *P. swiftii.* Phyogenetic analysis of these markers showed the two symbionts were extremely closely related: across the nuclear 18S and 28S rRNA genes (5054 bp) they shared 99.4% identity, and across the three mitochondrial genes (2984 bp) they shared 99.5% identity. We examined the rest of the transcriptome data to determine if this high level of identity was due to cross-contamination, and found the presence of zoanthid (host contaminant) sequences only in the *P. swiftii* sample and Symbiodiniaceae reads only in the sample from *M. mirabilis* (a photosynthetic coral). This indicates that our results are authentic and that the two hosts indeed contain very closely related symbionts.

Together with the existing sequence data from *C. aquarius* (Kwong et al. 2019), we generated phylogenetic trees of both nuclear 18S + 28S rRNA and mitochondrial proteins (Fig. 2), and find that the corallicolids form a distinct, strongly supported clade that is separate from all other recognized apicomplexan taxa (Adl et al. 2019). Accordingly, we formally propose Corallicolida ord. nov., Corallicolidae fam. nov., to encompass the whole lineage of corallicolids.

**Figure 2.**
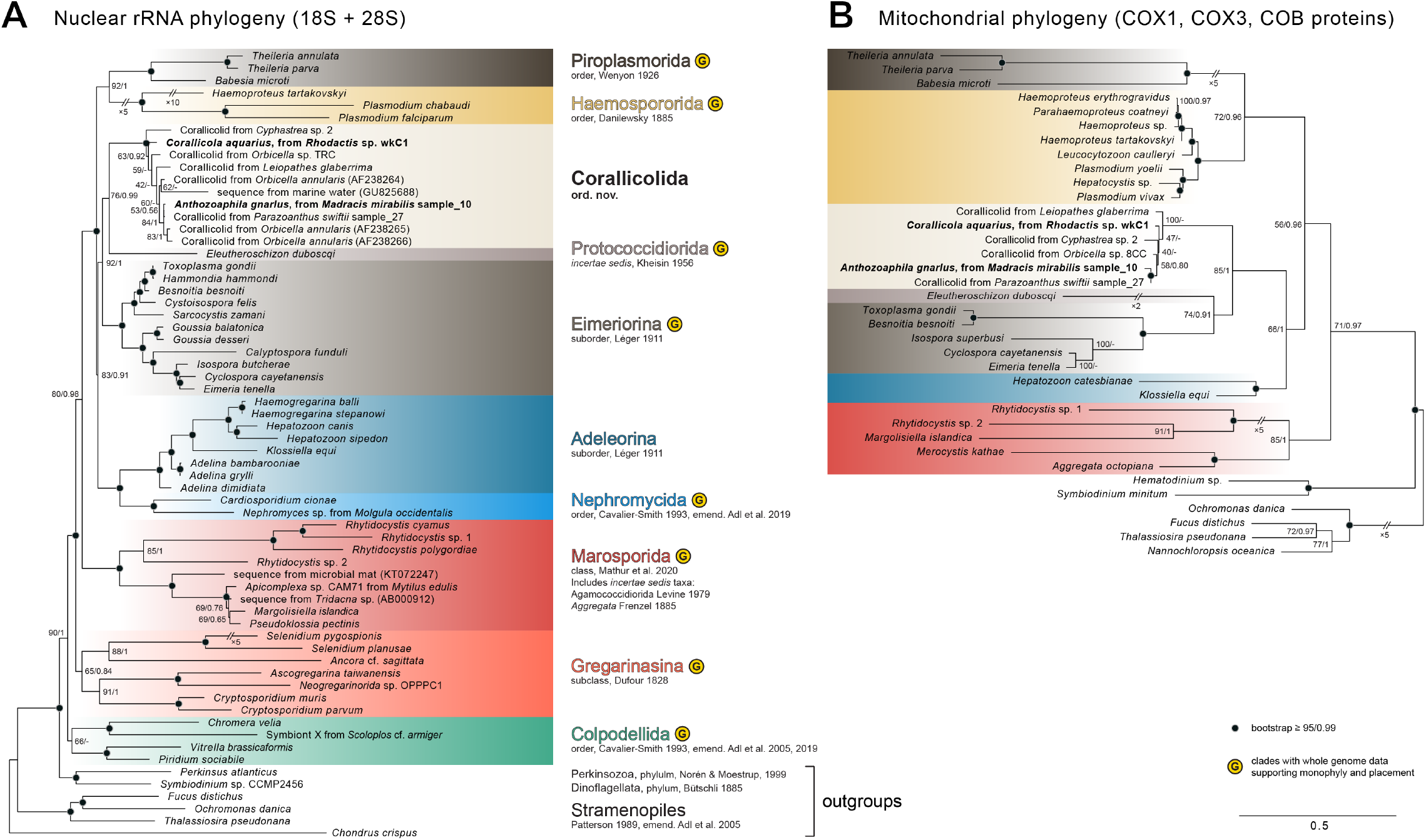
Phylogenetic placement of order Corallicolida. **A.** Nuclear rRNA gene phylogeny of the major divisions within the Apicomplexa. Tree was built with the maximum likelihood algorithm, using the TIM2+R5 substitution model. Clades supported by phylogenomic studies are marked. **B.** Mitochondrial protein phylogeny of the Apicomplexa. Clades are colored as in **A.** Tree was built with the maximum likelihood algorithm, using the LG+F+I+G4 substitution model. Bootstrap support percentage indicated at nodes. Posterior probabilities from Bayesian tree, built using the **A.** GTR+G and **B.** LG+I+G models, indicated after the slash. Bar represents average substitutions per site, with same scale for both trees.

Both nuclear and mitochondrial data suggest that the Corallicolida are related to coccidians, and probably most closely to Eimeriorina (e.g., *Toxoplasma*, *Eimeria*) and Protococcidiorida to the exclusion of Adelorina. This is consistent with previous analyses based on 18S rRNA data (Toller et al. 2012, del Campo et al. 2019), but contrasts with the assignment of *G. cylindrus* to the Agamococcidiorida based on morphological analysis (Upton and Peters 1986), or the more basally branching position within the Apicomplexa as suggested by plastid phylogenies (Janouškovec et al. 2012). The resolution of these questions will require additional genomic data from representatives of both Corallicolida and Adeleorina, but all extant data are in agreement that Corallicolida do not branch within any of the recognized apicomplexan lineages.

Phylogenetic inference for apicomplexans based on plastid data appears to be particularly problematic. It has recently been shown that plastid gene trees are incongruent with with phyogenomic trees based on nuclear data (which are considered to be much more signal-rich and, thus, accurate) (Muñoz-Gómez et al. 2019; Mathur et al. 2020). But plastid 16S rRNA data remains useful for assessing fine-scale diversity within this clade, because the density of sampling of corallicolids is significantly greater than with any other marker. An alignment of all available corallicolid 16S rRNA sequences longer than 1000 bp showed an average of 98.6% identity between sequences, with phylogenetic clustering largely based on host origin (Fig. S2). These results recapitulate previous findings based on short 16S rRNA fragments that uncovered corallicolid variation between coral host species, as well as between ecological habitats (Kwong et al. 2019; Vohsen et al. 2020). Molecular data consistently show a clear separation between *C. aquarius* and *A. gnarlus* (16S rRNA at 98.6% identity, 18S rRNA at 97.3% identity based on trimmed, near-full length alignments), which supports splitting these taxa into different genera. Other divergent sequences, such as that from the sea fan *Gorgonia ventalina* and that from a corallicolid plastid genome sequenced from the deep-water black coral *Leiopathes glaberrima* (Vohsen et al. 2020), may represent other species of corallicolids (Fig. S2B). A few sequences of apicomplexans that may be related to corallicolids, but in non-coral hosts, have been reported, such as a 16S rRNA sequence from anchovy gut (Fig. S2B), and the 18S rRNA sequences of some fish blood parasites (Hayes and Smit 2019). How the organisms bearing these sequences relate to Corallicolida remain to be examined.

Recognizing corallicolid diversity at the level of genera also provides more flexibility for reconciliation of this classification with *G. cylindrus*, which is a difficult problem. There is currently no molecular evidence that *G. cylindrus* is a member of this clade, but similarities in morphology and localization within coral mesentarial filaments suggest that it is a corallicolid. Arguably, given the nature of its description and the range of hosts attributed to it (it was described from several coral species, including its type host *D. cylindrus*), it may or may not represent a single discrete species. Molecular data from *D. cylindrus,* and its other reported hosts (Upton and Peters 1986), are required to clear up these issues and to prove that *G. cylindrus* shares genetic traits consistent with corallicolids. Nonetheless, it is apparent based on current evidence that its assignment to the Agamococcidiorida is incompatible with its membership with the Corallicolida. As such, we propose reassigning *G. cylindrus* to the Corallicolida, but keeping the designation of its distinct genus until further evidence can show where in the corallicolid phylogeny this taxon falls. By erecting the Corallicolida, and with the molecular and morphological identification of two species, *C. aquarius* and *A. gnarlus*, we provide a foundation for investigating of the diversity and distribution of this group.

### Future considerations

Corallicollids have now been found in a wide variety of anthozoan hosts (of which there are thought to be over 5,500 species [Crowther 2011]). They show a degree of host specificity, and molecular surveys consistently reveal substantial sequence-level diversity as well, altogether indicating that many new species and genera exist within the Corallicolida. Intriguingly, our analysis of corallicolid single cells from two anthozoans from Curaçao, the yellow pencil coral *M. mirabilis* and the sponge-dwelling zoanthid *P. swiftii*, suggested that similar symbionts strains can be found in different anthozoan hosts. This raises the prospect that some corallicolid lineages may live as generalists, while others may be more host restricted (Voshen et al. 2020). Further work will hopefully shed light on the morphological, behavioral, and genomic variation in the corallicolid symbionts of divergent coral hosts, as well as better define their life cycles and host ranges.

Closer scrutiny of the life cycle of corallicolids will additionally help clarify the nature of their interaction with coral hosts. Some apicomplexans have complex life cycles with alternating host species, but there is currently no evidence that this is the case with Corallicolida, as their sequences have not been detected in any host else besides anthozoans (Mathur et al. 2018; Kwong et al. 2019). This suggests that they may be monoxenous. We observed both intracellular and motile forms of corallicolids, and sporadic oocyst-like structures were also observed (not shown), the identity of which remains to be genetically verified.

Lastly, the overall position of corallicolids within the Apicomplexa as a whole is becoming more clear, but still needs to be confirmed by phylogenomics. The outcome will affect how virtually every characteristic described in the future will be interpreted. For example, the impact of their retention of the ancestral chlorophyll biosynthetic genes (Kwong et al. 2019) on how we reconstruct the loss of photosynthesis would be very different if we accepted the basally-branching position in plastid phylogenetic trees versus the coccidian-related position in nuclear and mitochondrial trees. By formalizing the classification of Corallicolida, a pivotal clade with ‘transitional’ features, we can begin tackling these evolutionary questions with renewed taxonomic clarity.

## TAXONOMIC SUMMARY

Phylum Apicomplexa Levine 1980, emend. Adl et al. 2005

Class Conoidasida Levine 1988

Order Corallicolida Kwong et al. 2020

Family Corallicolidae Kwong et al. 2020

Genus *Corallicola* Kwong et al. 2020

Species *Corallicola aquarius* Kwong et al. 2020

Genus *Anthozoaphila* Kwong et al. 2020

Species *Anthozoaphila gnarlus* Kwong et al. 2020

Genus *Gemmocystis* Upton & Peters 1986

Species *Gemmocystis cylindrus* Upton & Peters 1986

### Corallicolida ord. nov. Kwong et al. 2020

#### Diagnosis

A distinct and well-supported phylogenetic clade within the Apicomplexa based on 18S rRNA gene sequences. Presence of an intracellular stage as well as extracellular trophozoites. Cells colorless with smooth surface. Spherical or ovoid nucleus visible under light microscopy, located centrally or towards the posterior. Refractile bodies are absent. Plastid genome encodes genes for chlorophyll biosynthesis. All members of the order have so far been exclusively found in association with Anthozoa.

### Corallicolidae fam. nov. Kwong et al. 2020

#### Included taxa

*Corallicola gnarlus* gen. nov. sp. nov., *Anthozoaphila gnarlus* gen. nov. sp. nov.

#### Diagnosis

so far as for Corallicolida ord. nov.

#### Type genus

*Corallicola* gen. nov.

#### Zoobank ID

92E743FA-86E9-41B2-9DCE-B7DEE47A4E18

### *Corallicola* gen. nov. Kwong et al. 2020

#### Etymology

L. n. *corallium* coral; L. suff. *-cola* from L. n. *incola* inhabitant, dweller; N.L. n.

(nominative in apposition) *Corallicola* coral-dweller.

#### Diagnosis

Forms a distinct and well-supported phylogenetic lineage within the Corallicolidae based on 18S rRNA gene sequences.

#### Type species

*Corallicola aquarius* sp. nov.

#### Zoobank ID

67BDC122-7134-42B1-9F96-B813FEF25CAC

### *Corallicola aquarius* sp. nov. Kwong et al. 2020

#### Etymology

L. mas. adj. *aquarius* pertaining to water, referring to its presence in a corallimorpharian from an aquarium.

#### Diagnosis

Trophozoites elongate with an approximate length and width of 12 μm and 3.5 μm, respectively. Trophozoites elongate, exhibiting gliding motility. Cell displays a unique rRNA sequence phylotype represented by the type sequence. Found within the mesenterial filaments of *Rhodactis* sp. host.

#### Type sequence

18S rRNA gene sequence (MH304760) and 28S rRNA gene sequence (MH304761).

#### Type habitat

Marine; host associated.

#### Type host

*Rhodactis* sp. Milne, Edwards & Haime, 1851 (Metazoa; Cnidaria; Anthozoa; Corallimorpharia; Discosomidae).

#### Type material

Fixed specimens have been deposited in the Beaty Biodiversity Museum at the University of British Columbia under accession number MI-PR150.

#### Zoobank ID

59AC9E6A-31E8-4E6A-B8F9-F6BCA5EB0DF9

### *Anthozoaphila* gen. nov. Kwong et al. 2020

#### Etymology

N.L. n. *Anthozoa* reference to class Anthozoa; N.L. adj. *phila* from Gr. adj. *phílos* friendly to, loving; N.L. n. *Anthozoaphila* loving Anthozoa.

#### Diagnosis

Forms a distinct and well-supported phylogenetic lineage within the Corallicolidae based on 18S rRNA gene sequences. The genus type species was isolated from Curaçao and possessed 97.3% identity in the 18S rRNA gene sequence to the type species of *Corallicola* gen. nov.

#### Type species

*Corallicola gnarlus* sp. nov.

#### Zoobank ID

842F200E-B18A-4B79-8183-E0C023B63B13

### *Anthozoaphila gnarlus* sp. nov. Kwong et al. 2020

#### Etymology

N.L. mas. adj. *gnarlus*, derived from Eng. adj. *gnarly*, referring to the challenging identification of this taxa, and to acknowledge previous work on coral-associated apicomplexans (*<u>G</u>emmocystis*, genotype <u>N</u>, <u>ARL</u>-V).

#### Diagnosis

Trophozoites elongate with an approximate length and width of 18 μm and 3 μm, respectively. Trophozoites elongate, exhibiting gliding motility. Cell displays a unique rRNA sequence phylotype represented by the type sequence. Associated with the host *Madracis mirabilis* in the southern Caribbean.

#### Type sequence

18S rRNA gene sequence (MW192638) and 28S rRNA gene sequence (MW192641).

#### Type locality

Curaçao (12.1217, −68.9700).

#### Type habitat

Marine; host associated.

#### Type host

*Madracis mirabilis* Duchassaing & Michelotti, 1860 (Metazoa; Cnidaria; Anthozoa; Scleractinia; Pocilloporidae).

#### Type material

Fixed specimens have been deposited in the Beaty Biodiversity Museum at the University of British Columbia under accession number MI-PR151.

#### Zoobank ID

D8A2C974-EC45-470A-B6D7-402CB5241D44

## AUTHOR CONTRIBUTIONS

W.K.K. and P.J.K. conceived the study. W.K.K., N.A.T.I., N.O., M.J.A.V., and P.J.K. performed research. W.K.K., V.M., and I.N. analyzed data. W.K.K. and P.J.K. wrote the manuscript, with input from all authors.

## ACKNOWLEDGEMENTS

We thank the CARMABI research station staff for facilities access and technical assistance, Racquel Singh and Emma George for help in sample collection, and Kevin Wakeman for advice on cell isolation. W.K.K. was supported by the Natural Sciences and Engineering Research Council of Canada (NSERC) Fellowship PDF-502457-2017 and a Killam Postdoctoral Research Fellowship, N.A.T.I. was supported by an NSERC Canadian Graduate Scholarship, and V.M. was supported by a graduate fellowship from the University of British Columbia.

## SUPPLEMENTARY DATA

**Figure S1.** Light micrographs of corallicolid cell isolates used for single cell RNA-seq. **A.** Sample_10 from *M. mirabilis*. **B–F.** The five cells pooled to form sample_27 from *P. swiftii*. Scale bar = 5 μm, applicable to all panels. Note that cell morphology changes upon manipulation and washings used in the isolation protocol, when these images were taken, and are not necessarily representative of live motile cells (which are typically more elongated).

**Figure S2.** Diversity within Corallicolida. **A.** Plastid rRNA gene phylogeny of corallicolids and related taxa. Sequences from rRNA sequence surveys (i.e., not from isolates or plastid genome assemblies) are colored according to host origin. Dots at nodes indicate ≥ 95% bootstrap support. Tree was built with the maximum likelihood algorithm, using the GTR+R3 substitution model. **B.** Haplotype network of the corallicolid 16S rRNA gene (1419 bp trimmed length). Each hash mark indicates a difference at a single site. Size of nodes corresponds number of sequences. To indicate host of origin, nodes are colored as in **A.** Intermediate nodes are indicated by small black dots.

**Supplementary Table 1:** List of sequences used in building phylogenetic trees.

**Supplementary Videos 1 & 2:** Gliding motility of *A. gnarlus* cell from crushed *M. mirabilis* slurry.

