## Supplementary figures and images for "Taxonomy of the apicomplexan symbionts of coral, including Corallicolida ord. nov., reassignment of the genus *Gemmocystis,* and description of new species *Corallicola aquarius* gen. nov. sp. nov. and *Anthozoaphila gnarlus* gen. nov. sp. nov"

### Figure S1

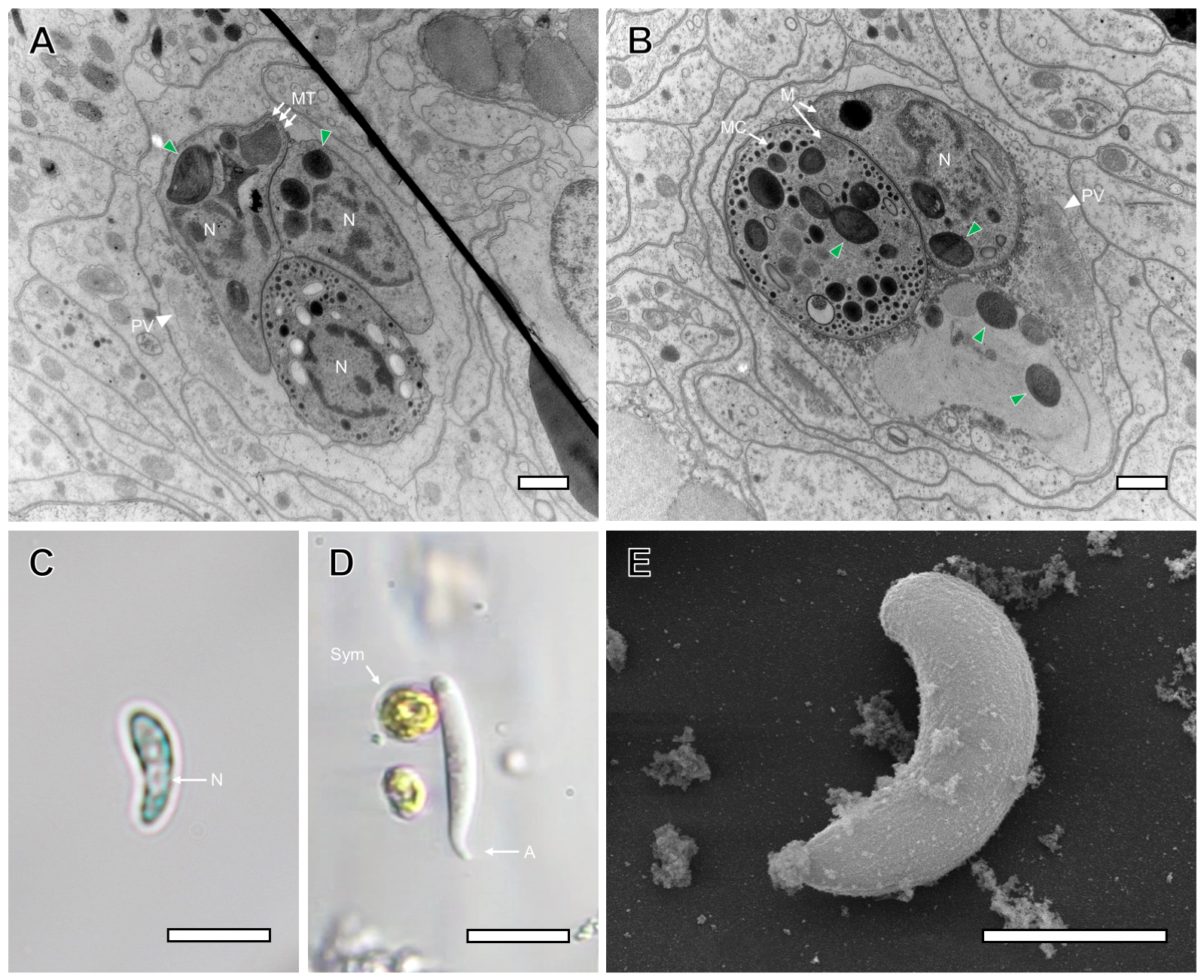

### Figure S2

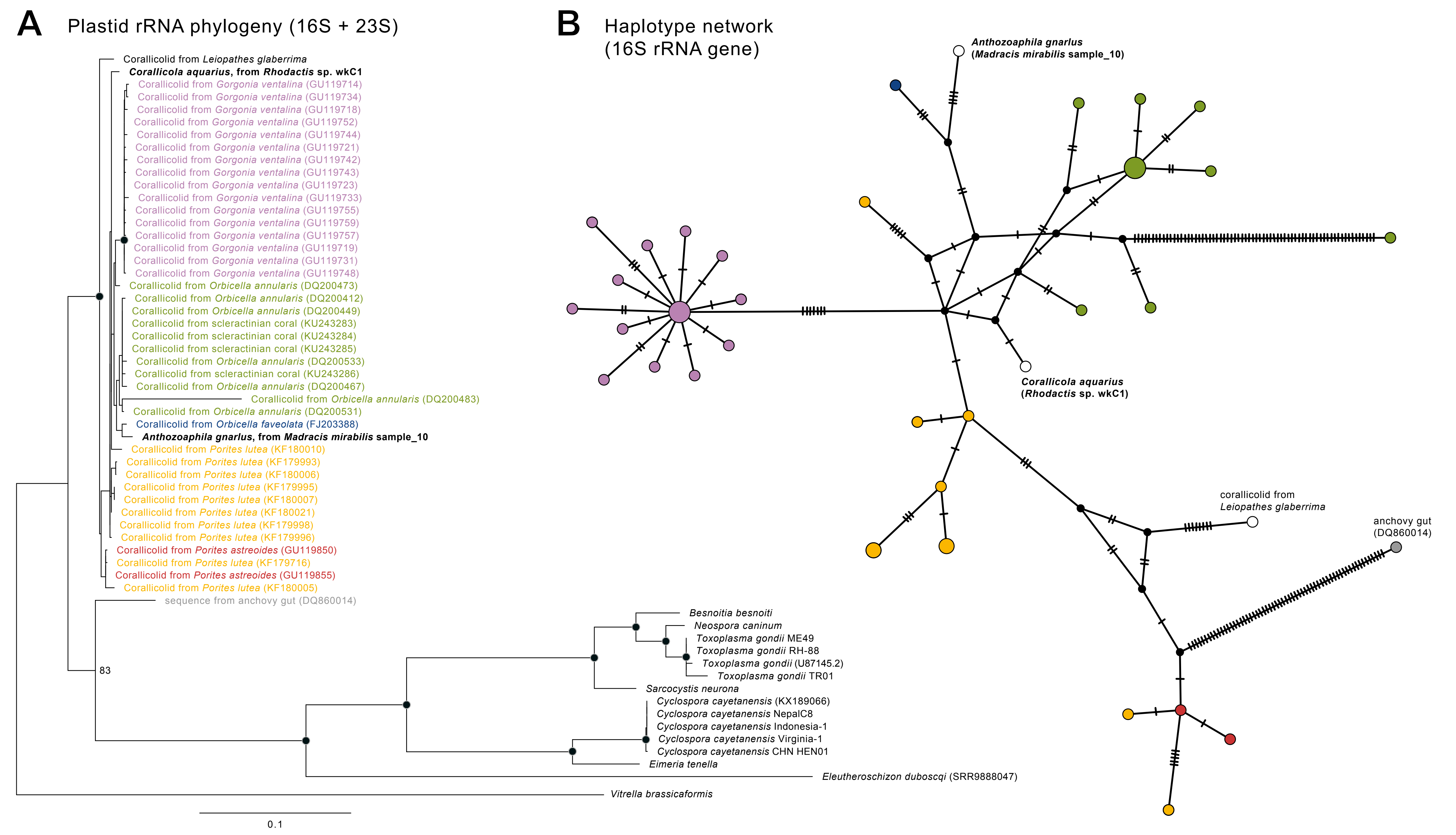
